# The global carrier frequency and genetic prevalence of Upshaw-Schulman syndrome

**DOI:** 10.1101/2021.02.28.433213

**Authors:** Ting Zhao, Shanghua Fan, Liu Sun

## Abstract

**Purpose:** Upshaw–Schulman syndrome (USS) is an autosomal recessive disease of thrombotic microangiopathy, caused by pathogenic variants in ADAMTS13. We aimed to (1) perform data mining pathogenicity of ADAMTS13 variants, (2) estimate carrier frequency and genetic prevalence of USS from gnomAD data, and (3) curated ADAMTS13 gene pathogenic variants dataset.

**Methods:** PubMed and Scopus were comprehensive retrieved. All previously reported pathogenic ADAMTS13 variants were compiled and annotated with gnomAD allele frequencies. Pooled global and population-specific carrier frequency and genetic prevalence for USS were calculated using Hardy-Weinberg equation.

**Results:** we mined reported disease-causing variants, of these were present in gnomAD exomes v2.1.1, filtering by allele frequency, pathogenicity of variants were classified by American College of Medical Genetics and Genomics criteria. The genetic prevalence and carrier frequency of USS was 0.43 per 1 million (95% CI: [0.36, 0.55]) and 1.31 per thousand, respectively. Combining known with novel pathogenic/likely pathogenic variants, the genetic prevalence and carrier frequency are 1.1 per 1 million (95% CI: [0.89, 1.37]) and 2.1 per thousand, respectively.

**Conclusion:** the genetic prevalence and carrier frequency of Upshaw-Schulman syndrome are within range of previously rough estimated.

## Introduction

Upshaw–Schulman syndrome (USS), also known as hereditary/congenital/familial thrombotic thrombocytopenic purpura (TTP), is an ultra-rare but life-threatening autosomal recessive disease that is absence or severe deficiency of plasma von Willebrand factor (vWF)-cleaving protease ADAM metallopeptidase with thrombospondin type 1 motif 13 (ADAMTS13), resulting in the abnormal presence of ultra-large vWF multimers and subsequent platelet adhere to vWF multimers leading to formation of circulating platelet microthrombi.^1-3^ The spectrum of clinical phenotype in USS is wide. Disease onset ranged from neonatal, childhood, adult to elder with a notable peak in women during pregnancy. The manifestations as recurrent attacks of microvascular thrombosis with associated thrombocytopenia, purpura and microangiopathic hemolytic anemia (MAHA), which leads to the ischemic damage of end organs (kidney, heart, and brain). Diagnosis is based on pentad of classic clinical characteristics: thrombocytopenia, hemolytic anemia, renal failure, fever, and neurologic deficits.^4,5^ Combined ADAMTS13 activity assay with genetic testing distinguishes USS from acquired TTP. Treatment in USS involves replacement of ADAMTS13 by fresh-frozen plasma (FFP) infusion.

USS is the result of homozygous or compound heterozygous variants in ADAMTS13 gene. ADAMTS13 gene spans 29 exons and ∼37 kb and is located at chromosome 9q34 and encodes a protein with 1,427 amino acids.^6^ to data, more than 200 ADAMTS13 disease-causing mutations, spread over all ADAMTS13 exons, have been identified in patients with USS since 2001.^7-12^

USS is extremely rare, and its precise prevalence is uncertain. Although most estimates suggest a prevalence of 0.5 to 2 cases per million population. The previously only two reported prevalence of Upshaw-Schulman syndrome were extremely heterogeneous, in central Norway was 16.7 per 1 million whereas in Norway was 3.1 per 1 million,^13^ which was 18 times / 3.4 times higher than the prevalence of USS in Japan (1 per 1.1 million),^14^ respectively. We guessed that the prevalence of USS will vary among different population or ethnicities.

Therefore, we aimed to estimate the prevalence of USS across ethnicities from current largest public available gnomAD exomes dataset using validated protocols.^15,16^ in addition, we aim to generate evidence-based dataset of known USS pathogenic variants by data mining. Meanwhile, we also aim to generate machine learning training dataset for pathogenicity interpretation of variants.

## MATERIALS AND METHODS

### Identification of known disease-causing variants

A comprehensive literature retrieval of PubMed and Scopus was performed to identify all known disease-causing variants in the ADAMTS13 gene (see supplementary materials for search terms, protocols, scripts, full paper list and full variants list).

Two independent reviewers reviewed titles and abstracts to screen articles according to inclusion and exclusion criteria: original case reports reported disease-causing variants within the ADAMTS13 gene were included, variants in full-text tables, figures or supplementary material figures and tables were extracted. non-English articles, reviews, comments, and editorials *etc*. nonoriginal papers and *in vitro* or animal model studies were excluded.

All paper were saved in Medline format and stored in NoSQL database MongoDB documents using NCBI Entrez Programming Utilities^17^ (E-utilities) with Python package biopython^18^ and pymongo implementation.

The database HGMD^19^ (http://www.hgmd.cf.ac.uk/ac/index.php), Ensembl Variation^20^, VarSome^21^ (https://varsome.com/), ClinVar^22^ (https://www.ncbi.nlm.nih.gov/clinvar/) and Genomenon Mastermind^23^ (https://mastermind.genomenon.com/) were used alongside literature retrieval to identify additional ADAMTS13 variants with published reports of pathogenicity.

A list of all single nucleotide variants (SNVs) for ADAMTS13 was compiled using Ensembl Variant Simulator.^24^

### Identification of major functional variants

Genome Aggregation Database (gnomAD) ^25^ was searched for pathogenic variants that have not yet been reported in patients, we examined all caused major functional or structural changes (frameshift, stop codon, initiation codon, splice donor and splice acceptor).

### Annotation of variants with allele frequency and functional predictions

Raw variants were identified and converted to Human Genome Variation Society (HGVS) nomenclature^26^ using Mutalyzer^27^ and Ensembl VEP Variant Recoder REST API with Python implementation. Ensembl variant effect predictor (VEP)^28^ was used to annotate variants and *in silico* predictions of pathogenicity with PROVEAN/PolyPhen/MutationTaster. gnomAD minor allele frequency (MAF) data was added to each variant from gnomAD browser.

### Disease-causing variants classification

Pathogenicity of variants were interpreted using a pipeline proposed by Zhang et al.^29^ Disease-causing variants with gnomAD allele frequency were classified using the American College of Medical Genetics and Genomics (ACMG) and the Association for Molecular Pathology (AMP) criteria^30^ with ClinGen Pathogenicity Calculator^31^. Pathogenic/likely pathogenic variants were included for prevalence calculation.

### Maximum population filtering allele frequency

All variants with gnomAD allele frequency data were filtered using a method defined by Whiffin et al.^32^ Prevalence was taken from estimate from Japanese^14^ population and orphanet database: one case per 1 million. Maximum allelic contribution was set at 24.4% based on estimate of c.4143dup (p.Glu1382Argfs*6) from the International Hereditary Thrombotic Thrombocytopenic Purpura Registry^7^ data. Maximum genetic contribution set on 1 based on UK^8^, French^9^, Germany^10^ cohorts and the International Hereditary Thrombotic Thrombocytopenic Purpura Registry^7^ data. The penetrance was set as 50%, which was suggested by Whiffin’s methods. The maximum population credible allele frequency was calculated as 0.035% by Whiffin’s defined equation.

The maximum population filtering allele frequencies were directly downloaded from gnomAD browser (https://gnomad.broadinstitute.org/). Variants with maximum population filtering allele frequency greater than the maximum population credible allele frequency were excluded.

### Prevalence calculation

Allele frequencies of pathogenic/likely pathogenic variants were collected from ADAMTS13 variant dataset, pooled, and calculated of the prevalence of USS using the Hardy-Weinberg equation.

Binomial proportion 95% confidence interval (95% CI) were calculated using Wilson score with Python scientific computing package statsmodels and NumPy implementation. Graphics were plotted using R package ggplot2 and VennDiagram^33^.

## RESULTS

### Identification of *ADAMTS13* variants

Comprehensive retrieval for USS disease-causing variants returned 1249 articles, of which 126 literatures were included according to exclusion and inclusion criteria, from which 280 disease-causing variants were collected, of which 239 variants were classified as “pathogenic” or “likely pathogenic” under ACMG criteria. Retrieving ClinVar database return additional 6 disease-causing variants (pathogenic and likely pathogenic). A total of 245 known disease-causing variants were mined. gnomAD allele frequencies were available for 59/245 (24.1%) of disease-causing variants. All disease-causing variants pipeline and count in Figure 1 and associated data see supplementary data.

**Fig. 1.**
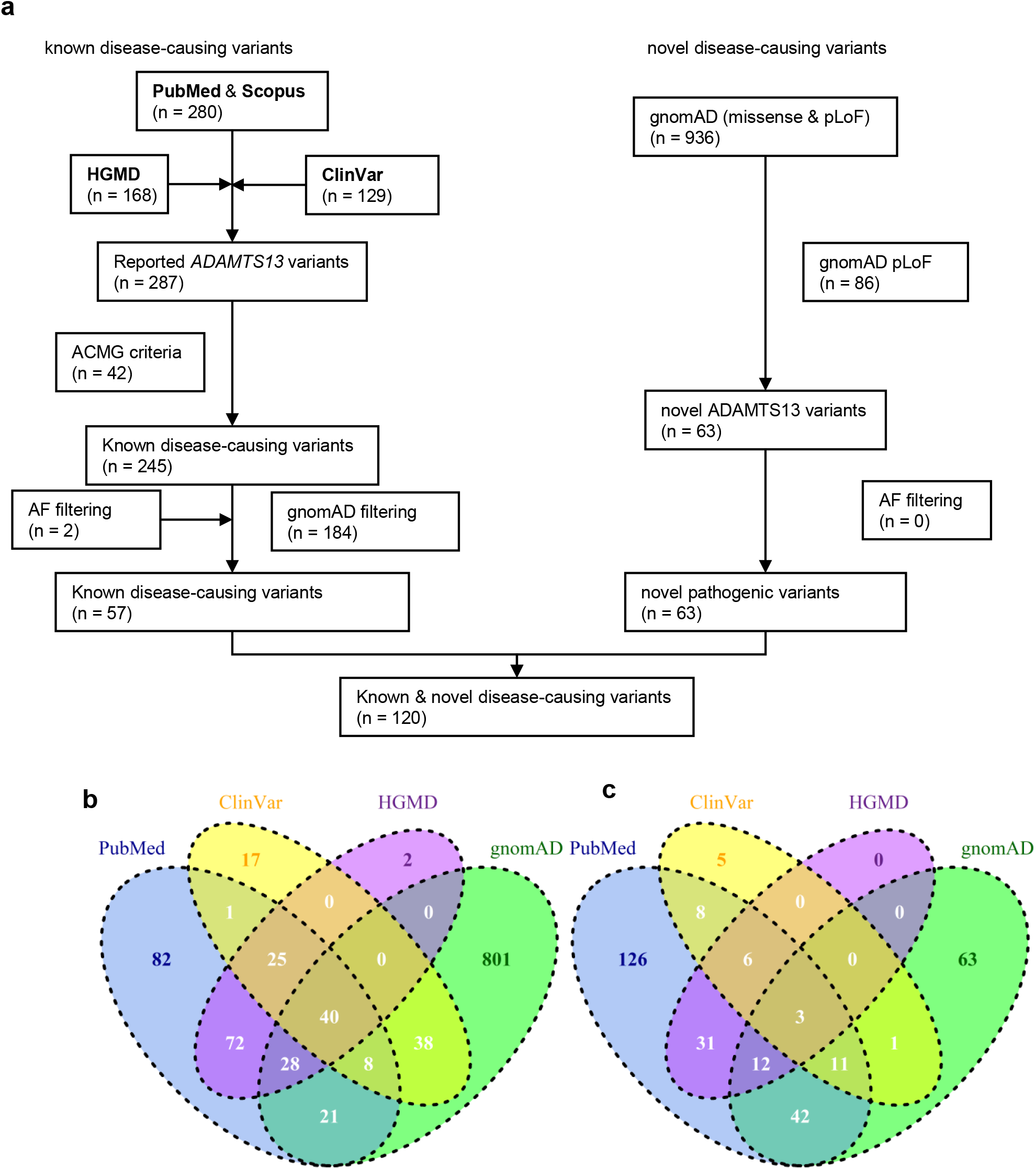
*ADAMTS13* gene disease-causing variants and gnomAD allele frequencies. (**a**) flow chart of identification and classification of *ADAMTS13* disease-causing variants. *ADAMTS13* variants were extracted from PubMed & Scopus citations. *ADAMTS13* missense, nonsense, frameshift, inframe, splice acceptor / donor variants were collected from HGMD Public (2016 version), ClinVar and gnomAD database. (**b**) Venn diagram of mined PubMed & Scopus, HGMD, ClinVar and gnomAD variants. (**c**) Venn diagram of mined PubMed & Scopus, HGMD, ClinVar and gnomAD disease-causing variants.

### Frequency of reported USS pathogenic/likely pathogenic variants

Of 59 reported disease-causing variants with gnomAD allele frequency data, 57 remained after frequency filtering. Pooling of the allele frequencies of these variants calculate a global allele frequency of 0.0006, which is equivalent to prevalence of 0.43 per 1,000,000 (95% confidence interval: [0.36, 0.55]). five major populations had similar prevalence of less than 1 per million (Figure 2 and Table 1).

**Table 1.** allele frequency data-based prevalence and carrier frequency calculation.

|  | prevalence |  | carrier frequency |  |
| --- | --- | --- | --- | --- |
|  | known and novel variants | known variants | known and novel variants | known variants |
| **total** | 1.10152<br>(0.890567, 1.370326) | 0.428407<br>(0.3357, 0.554897) | 0.002097 | 0.001308 |
| **AFR** | 5.639105<br>(3.010004, 10.55961) | 0.944298<br>(0.355737, 2.505441) | 0.004738 | 0.001942 |
| **AMR** | 1.482111<br>(0.812177, 2.704048) | 0.565474<br>(0.263468, 1.213389) | 0.002432 | 0.001503 |
| **ASJ** | 0.046311<br>(0.003816, 0.561623) | 0 | 0.00043 | 0 |
| **EAS** | 1.676507<br>(0.755126, 3.720573) | 0.781969<br>(0.298701, 2.046257) | 0.002586 | 0.001767 |
| **FIN** | 0.00864<br>(0.000654, 0.114046) | 0 | 0.000186 | 0 |
| **NFE** | 1.143383<br>(0.595458, 1.961721) | 0.593037<br>(0.239053, 1.177138) | 0.002136 | 0.001539 |
| **SAS** | 1.121036<br>(0.56517, 2.22306) | 0.436036<br>(0.183617, 1.035195) | 0.002115 | 0.00132 |
| **OTH** | 0.731709<br>(0.138862, 3.850784) | 0.107322<br>(0.008125, 1.415878) | 0.001709 | 0.000655 |

AFR: African/African American, AMR: Latino/Admixed American, ASJ: Ashkenazi Jewish, EAS: East Asian, FIN: Finnish, NFE: Non-Finnish European, SAS: South Asian, OTH: Other.

**Fig. 2.**
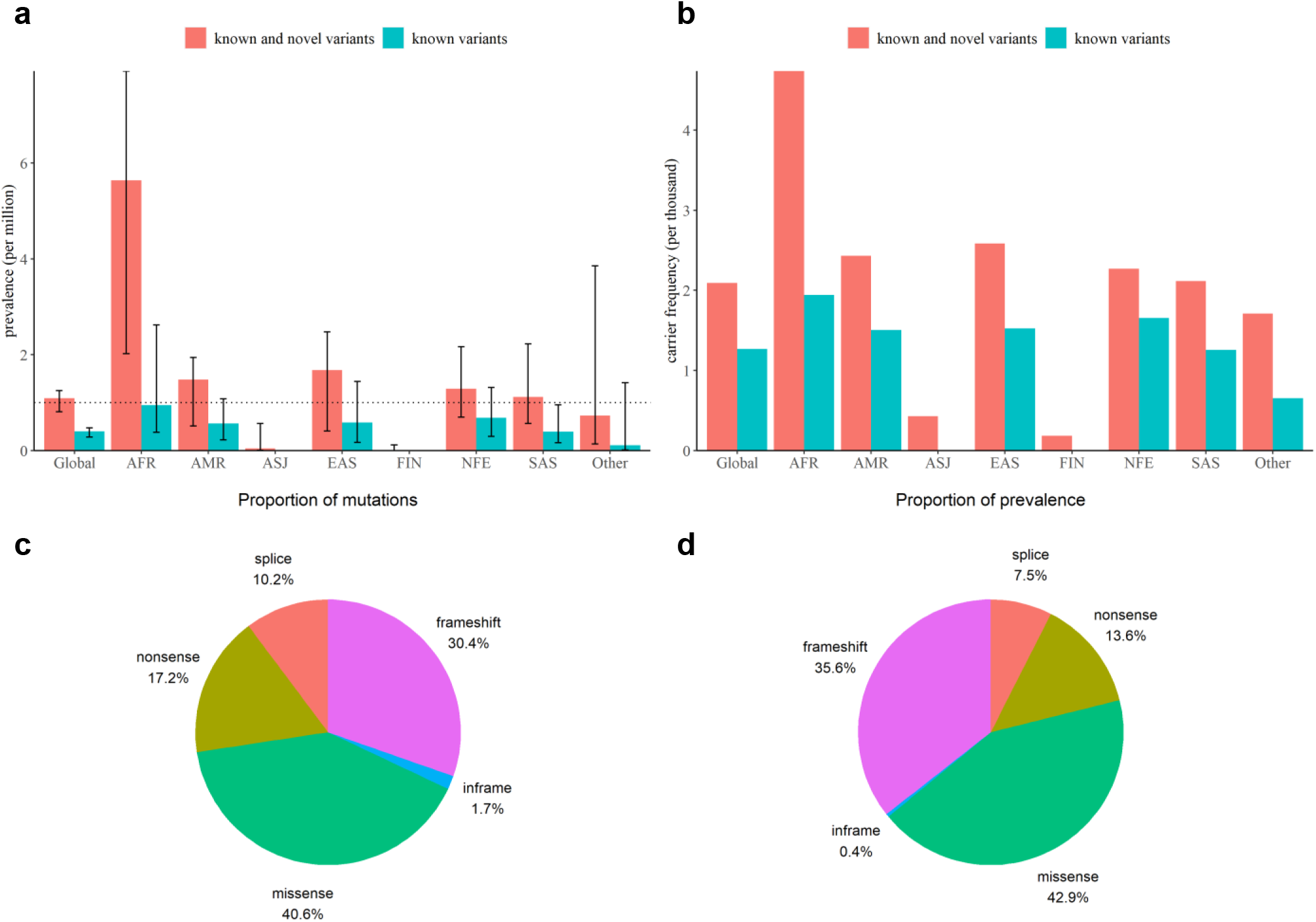
genetic prevalence and carrier frequency of USS. (**a, b**) USS carrier frequency and genetic prevalence estimated from gnomAD allele frequencies. (**c, d**) molecular consequence of all known and novel disease-causing variants. (c) Pie chart of the number of variants group by each molecular consequence. (d) Pie chart of the proportion of the total allele frequency group by molecular consequence.

### Functional pathogenic variants

To estimate the genetic prevalence of USS, including disease-causing variants that have not yet been reported in patients, we searched all ADAMTS13 variants in gnomAD database that caused loss of function (LoF) mutation (frameshift, nonsense, splice acceptor and splice donor variants). After filtering, 86 variants were found in gnomAD exomes v2.1.1 database and 63 variants were novel. combined novel disease-causing variants with reported pathogenic variants, the global allele frequency of USS was 0.001, Equivalent to 1.1 per 1,000,000 (95% confidence interval: [0.89, 1.37]). The African population had the highest prevalence of 5.64 per 1,000,000 (95% CI: [3.01, 10.56]) and the other four major population had prevalence of great than 1 per 1,000,000.

The most common mutation functional consequence was missense, accounting for 40.6% of all pathogenic and likely pathogenic variants and contributing 42.9% of the total allele frequency. Meanwhile, frameshift and nonsense were the second most common.

## DISCUSSION

In this study, we conducted the first-time systematic study to unbiased estimate the genetic prevalence of USS both global and five major populations. Our result was within the range previously rough estimated. Meanwhile, we have manual collected all ADAMTS13 disease-causing variants with evidence-based interpretation of pathogenicity.

USS accounts for <5% of TTP cases and is caused mostly by biallelic (compound heterozygote or homozygote) mutation in the ADAMTS13 gene, or in rare cases, by monoallelic ADAMTS13 mutation associated with single-nucleotide polymorphisms (SNPs). USS has heterogeneous inheritance pattern. Previous estimates of USS prevalence were heterogeneous, which may be largely accounted for by differences in population. Consequently, using the current largest population genome dataset, gnomAD v2.1.1 (125,748 human exomes and 15,708 genomes), we calculated that global genetic prevalence of USS is 0.43 to 1.1 per million and carrier frequency is 1 to 2 per thousand. We have highlighted African population have highest prevalence of USS, other four major population have similar prevalence and carrier frequency. USS was not in the first Rare Diseases List released by China goverment.^34^ The prevalence of USS in Chinese population was not ever estimated.^35^ we have demonstrated the power and limitations of population genome dataset to calculate genetic prevalence and carrier frequency of USS. gnomAD group East Asians population roughly into three categories: Korean, Japanese and other East Asians. Other population genome dataset, 100k Chinese people genome project and GenomeAsia 100K Project will fill the gap.^36^ We will estimate prevalence of Upshaw-Schulman syndorme in Asian population and Chinese population with 100k genome datasets in the next step.

Two variants, c.3178C>T (p.Arg1060Trp) and c.559G>C (p.Asp187His), which were classified pathogenic and likely pathogenic respectively, were filtered out by Whiffin’s method, which were “too common” to be causative for USS based on our set value for maximum allelic contribution and prevalence. Whiffin’s method was not optimal but more persuasive than arbitrary minor allele frequency (MAF) cutoff threshold 0.05 (ACMG benign stand-alone criteria).

We study based on assumptions of Hardy-Weinberg equation. But consanguine marriage is popular in specific subpopulation (such as Africa and south Asia). Genetic prevalence might be higher than calculated values. In addition, our genetic prevalence calculation algorithm is just one. Other algorithm, such as product-based algorithm for allele matrices and Bayesian-based algorithm calculated autosomal recessive inherited retinal diseases^37^ and limb-girdle muscular dystrophy^38^, respectively.

The number of ADAMTS13 classified variants in ClinVar database were far less than reported variants by document retrieval and data mining, but pathogenicity prediction tools, used ClinVar dataset as training set. Clinical Genome (ClinGen) can be used in variants evaluation and assertion. dbNSFP database, which comprehensive functional prediction and annotation for human nonsynonymous and splice-site SNVs, was valuable resource for training set construction in pathogenicity prediction of novel variants.^39^

Our finding of reported disease-causing variants and predicted pathogenic variants highlighted the mutational spectrum of USS. The most common pathogenic variants were missense, and which also were most difficult to predict and evaluate pathogenicity. it will lead to toolboxes for geneticists, clinicians, genetic counselors, and health data analysts.

In summary, the genetic prevalence of USS is like estimated, we have also calculated more reliable global and population-specific estimates for USS genetic prevalence and carrier frequency, these data can be used training set for pathogenicity prediction of novel variants and gene diagnosis of Upshaw-Schulman syndrome. We also provided validated pipeline to calculate prevalence of rare diseases. These datasets will be especially valuable for rare diseases definition in developing country which are scarcity of epidemiological data.^40^

## ACKNOWLEDGEMENTS

This study was supported by Yunnan Fundamental Research Projects (grant No. 202101AU070007).

## DATA REPOSITORY

Associated data will also be founded at Science Data Bank (ScienceDB) [DOI: 10.11922/sciencedb.00628]. Code can be founded at GitHub (https://github.com/liu-sun/USS).

## References

1. Kremer Hovinga JA, George JN. Hereditary Thrombotic Thrombocytopenic Purpura. N Engl J Med. 2019;381(17):1653–1662.

2. Kremer Hovinga JA, Coppo P, Lammle B, Moake JL, Miyata T, Vanhoorelbeke K. Thrombotic thrombocytopenic purpura. Nat Rev Dis Primers. 2017;3:17020.

3. Joly BS, Coppo P, Veyradier A. Thrombotic thrombocytopenic purpura. Blood. 2017;129(21):2836–2846.

4. Matsumoto M, Fujimura Y, Wada H, et al. Diagnostic and treatment guidelines for thrombotic thrombocytopenic purpura (TTP) 2017 in Japan. Int J Hematol. 2017;106(1):3–15.

5. Scully M, Hunt BJ, Benjamin S, et al. Guidelines on the diagnosis and management of thrombotic thrombocytopenic purpura and other thrombotic microangiopathies. Br J Haematol. 2012;158(3):323–335.

6. Levy GG, Nichols WC, Lian EC, et al. Mutations in a member of the ADAMTS gene family cause thrombotic thrombocytopenic purpura. Nature. 2001;413(6855):488–494.

7. van Dorland HA, Taleghani MM, Sakai K, et al. The International Hereditary Thrombotic Thrombocytopenic Purpura Registry: key findings at enrollment until 2017. Haematologica. 2019;104(10):2107–2115.

8. Alwan F, Vendramin C, Liesner R, et al. Characterization and treatment of congenital thrombotic thrombocytopenic purpura. Blood. 2019;133(15):1644–1651.

9. Joly BS, Boisseau P, Roose E, et al. ADAMTS13 Gene Mutations Influence ADAMTS13 Conformation and Disease Age-Onset in the French Cohort of Upshaw-Schulman Syndrome. Thromb Haemost. 2018;118(11):1902–1917.

10. Hassenpflug WA, Obser T, Bode J, et al. Genetic and Functional Characterization of ADAMTS13 Variants in a Patient Cohort with Upshaw-Schulman Syndrome Investigated in Germany. Thromb Haemost. 2018;118(4):709–722.

11. Miyata T, Kokame K, Matsumoto M, Fujimura Y. ADAMTS13 activity and genetic mutations in Japan. Hamostaseologie. 2013;33(2):131–137.

12. Fujimura Y, Matsumoto M, Isonishi A, et al. Natural history of Upshaw-Schulman syndrome based on ADAMTS13 gene analysis in Japan. J Thromb Haemost. 2011;9 Suppl 1:283–301.

13. von Krogh AS, Quist-Paulsen P, Waage A, et al. High prevalence of hereditary thrombotic thrombocytopenic purpura in central Norway: from clinical observation to evidence. J Thromb Haemost. 2016;14(1):73–82.

14. Kokame K, Kokubo Y, Miyata T. Polymorphisms and mutations of ADAMTS13 in the Japanese population and estimation of the number of patients with Upshaw-Schulman syndrome. J Thromb Haemost. 2011;9(8):1654–1656.

15. Gao J, Brackley S, Mann JP. The global prevalence of Wilson disease from next-generation sequencing data. Genet Med. 2019;21(5):1155–1163.

16. Wallace DF, Subramaniam VN. The global prevalence of HFE and non-HFE hemochromatosis estimated from analysis of next-generation sequencing data. Genet Med. 2016;18(6):618–626.

17. Sayers EW, Beck J, Bolton EE, et al. Database resources of the National Center for Biotechnology Information. Nucleic Acids Res. 2021;49(D1):D10–D17.

18. Cock PJ, Antao T, Chang JT, et al. Biopython: freely available Python tools for computational molecular biology and bioinformatics. Bioinformatics. 2009;25(11):1422–1423.

19. Stenson PD, Mort M, Ball EV, et al. The Human Gene Mutation Database (HGMD((R))): optimizing its use in a clinical diagnostic or research setting. Hum Genet. 2020;139(10):1197–1207.

20. Hunt SE, McLaren W, Gil L, et al. Ensembl variation resources. Database (Oxford). 2018;2018.

21. Kopanos C, Tsiolkas V, Kouris A, et al. VarSome: the human genomic variant search engine. Bioinformatics. 2019;35(11):1978–1980.

22. Landrum MJ, Chitipiralla S, Brown GR, et al. ClinVar: improvements to accessing data. Nucleic Acids Res. 2020;48(D1):D835–D844.

23. Chunn LM, Nefcy DC, Scouten RW, et al. Mastermind: A Comprehensive Genomic Association Search Engine for Empirical Evidence Curation and Genetic Variant Interpretation. Front Genet. 2020;11:577152.

24. Howe KL, Achuthan P, Allen J, et al. Ensembl 2021. Nucleic Acids Res. 2021;49(D1):D884–D891.

25. Karczewski KJ, Francioli LC, Tiao G, et al. The mutational constraint spectrum quantified from variation in 141,456 humans. Nature. 2020;581(7809):434–443.

26. den Dunnen JT, Dalgleish R, Maglott DR, et al. HGVS Recommendations for the Description of Sequence Variants: 2016 Update. Hum Mutat. 2016;37(6):564–569.

27. den Dunnen JT. Sequence Variant Descriptions: HGVS Nomenclature and Mutalyzer. Curr Protoc Hum Genet. 2016;90:7 13 11–17 13 19.

28. McLaren W, Gil L, Hunt SE, et al. The Ensembl Variant Effect Predictor. Genome Biol. 2016;17(1):122.

29. Zhang J, Yao Y, He H, Shen J. Clinical Interpretation of Sequence Variants. Curr Protoc Hum Genet. 2020;106(1):e98.

30. Richards S, Aziz N, Bale S, et al. Standards and guidelines for the interpretation of sequence variants: a joint consensus recommendation of the American College of Medical Genetics and Genomics and the Association for Molecular Pathology. Genet Med. 2015;17(5):405–424.

31. Patel RY, Shah N, Jackson AR, et al. ClinGen Pathogenicity Calculator: a configurable system for assessing pathogenicity of genetic variants. Genome Med. 2017;9(1):3.

32. Whiffin N, Minikel E, Walsh R, et al. Using high-resolution variant frequencies to empower clinical genome interpretation. Genet Med. 2017;19(10):1151–1158.

33. Chen H, Boutros PC. VennDiagram: a package for the generation of highly-customizable Venn and Euler diagrams in R. BMC Bioinformatics. 2011;12:35.

34. He J, Kang Q, Hu J, Song P, Jin C. China has officially released its first national list of rare diseases. Intractable Rare Dis Res. 2018;7(2):145–147.

35. He J, Tang M, Zhang X, et al. Incidence and prevalence of 121 rare diseases in China: Current status and challenges. Intractable Rare Dis Res. 2019;8(2):89–97.

36. GenomeAsia KC. The GenomeAsia 100K Project enables genetic discoveries across Asia. Nature. 2019;576(7785):106–111.

37. Hanany M, Rivolta C, Sharon D. Worldwide carrier frequency and genetic prevalence of autosomal recessive inherited retinal diseases. Proc Natl Acad Sci U S A. 2020;117(5):2710–2716.

38. Liu W, Pajusalu S, Lake NJ, et al. Estimating prevalence for limb-girdle muscular dystrophy based on public sequencing databases. Genet Med. 2019;21(11):2512–2520.

39. Liu X, Li C, Mou C, Dong Y, Tu Y. dbNSFP v4: a comprehensive database of transcript-specific functional predictions and annotations for human nonsynonymous and splice-site SNVs. Genome Med. 2020;12(1):103.

40. Haendel M, Vasilevsky N, Unni D, et al. How many rare diseases are there? Nat Rev Drug Discov. 2020;19(2):77–78.

