## supplementary data for "The global carrier frequency and genetic prevalence of Upshaw-Schulman syndrome"

Supplementary information

### Data mining Protocols

#### Literature retrieval

**PubMed** (<https://pubmed.ncbi.nlm.nih.gov/>)

search terms:

- “Purpura, Thrombotic Thrombocytopenic”[Mesh] “Mutation”[Mesh]
- Upshaw-Schulman syndrome[title]
- Congenital thrombotic thrombocytopenic purpura[title]
- Hereditary thrombotic thrombocytopenic purpura[title]
- Inherited thrombotic thrombocytopenic purpura[title]
- Familial thrombotic thrombocytopenic purpura[title]
- ADAMTS13[title] (mutation*[title] OR variant*[title] OR variation*[title])
- von Willebrand factor cleaving protease[title] AND (mutation*[title] OR variant*[title] OR variation*[title])

**Scopus** (<https://www.scopus.com/home.uri>)

Search terms:

(TITLE-ABS-KEY (“thrombotic thrombocytopenic purpura”) OR TITLE-ABS-KEY (“Upshaw-Schulman syndrome”)) AND (TITLE-ABS-KEY (mutation*) OR TITLE-ABS-KEY (variant*)) AND DOCTYPE (ar)

#### Database search

**ClinVar (**[**https://www.ncbi.nlm.nih.gov/clinvar/**](https://www.ncbi.nlm.nih.gov/clinvar/)**)**

ADAMTS13[gene] NOT (“copy number gain”[vartype] OR “copy number loss”[vartype] OR duplication[vartype])

### References

1.Manea, M. et al. ADAMTS13 phenotype in plasma from normal individuals and patients with thrombotic thrombocytopenic purpura. Eur J Pediatr 166, 249–257 (2007).

2.VEYRADIER, A., MEYER, D. & LOIRAT, C. Desmopressin, an unexpected link between nocturnal enuresis and inherited thrombotic thrombocytopenic purpura (Upshaw‐Schulman syndrome). J Thromb Haemost 4, 700–701 (2006).

3.Drews, K., Seremak-Mrozikiewicz, A., Sobieszczyk, S. & Barlik, M. Inherited thrombotic thrombocytopenic purpura in pregnancy. Neuro Endocrinol Lett 34, 508–13 (2013).

4.Lehmberg, K. et al. Inherited Thrombotic Thrombocytopenic Purpura (Upshaw Schulman Syndrome) as Differential Diagnosis to Neonatal Septicaemia with Disseminated Intravascular Coagulation ? a Case Series. Zeitschrift F R Geburtshilfe Und Neonatol 221, 39–42 (2016).

5.Richter, J. et al. Successful management of a planned pregnancy in severe congenital thrombotic thrombocytopaenic purpura: the Upshaw–Schulman syndrome. Transfusion Med 21, 211–213 (2011).

6.Sharma, D., Shastri, S., Pandita, A. & Sharma, P. Congenital thrombotic thrombocytopenic purpura: Upshaw–Schulman syndrome: a cause of neonatal death and review of literature. J Maternal-fetal Neonatal Medicine 29, 1977–1979 (2015).

7.Sakai, K. et al. Success and limitations of plasma treatment in pregnant women with congenital thrombotic thrombocytopenic purpura. J Thromb Haemost 18, 2929–2941 (2020).

8.Toret, E., Demir-Kolsuz, O., Ozdemir, Z. C. & Bor, O. A Case Report of Congenital Thrombotic Thrombocytopenic Purpura: The Peripheral Blood Smear Lights the Diagnosis. J Pediatric Hematology Oncol Publish Ahead of Print, 535–538 (2020).

9.Rashid, A., Mushtaq, N. & Mansoori, H. Congenital Thrombotic Thrombocytopenic Purpura With a Novel ADAMTS13 Gene Mutation. Cureus 12, e12053 (2020).

10.Vries, P. S. de et al. Genetic variants in the ADAMTS13 and SUPT3H genes are associated with ADAMTS13 activity. Blood 125, 3949–3955 (2015).

11.Chou, S.-C. et al. First reported case of congenital thrombotic thrombocytopenic purpura in Taiwan with novel mutation of ADAMTS13 gene. Int J Hematol 1–5 (2021) doi:10.1007/s12185-020-03068-5.

12.Edwards, N. C. et al. Characterization of Coding Synonymous and Non-Synonymous Variants in ADAMTS13 Using Ex Vivo and In Silico Approaches. Plos One 7, e38864 (2012).

13.Tsuda, M. et al. Upshaw-Schulman syndrome diagnosed during pregnancy complicated by reversible cerebral vasoconstriction syndrome. Transfus Apher Sci 57, 790–792 (2018).

14.Ferrari, B. & Peyvandi, F. How I treat thrombotic thrombocytopenic purpura in pregnancy. Blood 136, 2125–2132 (2020).

15.Xu, J., Yu, S. & Zhang, F. Frequent recurrence of pregnancy-triggered congenital thrombotic thrombocytopenic purpura: A rare case report. Clin Hemorheol Micro 1–6 (2020) doi:10.3233/ch-200970.

16.Kentouche, K. et al. Pregnancy in Upshaw-Schulman syndrome. H Mostaseologie 33, 144–148 (2013).

17.Borgi, A., Khemiri, M., Veyradier, A., Kazdaghli, K. & Barsaoui, S. Congenital Thrombotic Thrombocytopenic Purpura: Atypical Presentation And First ADAMTS 13 Mutation In A Tunisian Child. Mediterr J Hematology Infect Dis 5, 2013041 (2013).

18.Letzer, A. et al. Upshaw-Schulman syndrome-associated ADAMTS13 variants possess proteolytic activity at the surface of endothelial cells and in simulated circulation. Plos One 15, e0232637 (2020).

19.Manea, M. et al. Podocytes express ADAMTS13 in normal renal cortex and in patients with thrombotic thrombocytopenic purpura. Brit J Haematol 138, 651–662 (2007).

20.Morioka, M. et al. A first bout of thrombotic thrombocytopenic purpura triggered by herpes simplex infection in a 45-year-old nulliparous female with Upshaw-Schulman syndrome. Blood Transfus Trasfusione Del Sangue 12 Suppl 1, s153-5 (2013).

21.Fujimura, Y. et al. Patent ductus arteriosus generates neonatal hemolytic jaundice with thrombocytopenia in Upshaw-Schulman syndrome. Blood Adv 3, 3191–3195 (2019).

22.Bennett, M. et al. Experiences in a Family With the Upshaw-Schulman Syndrome Over a 44-Year Period. Clin Appl Thrombosis Hemostasis 20, 296–303 (2014).

23.Jiang, Y. et al. Novel mutations in ADAMTS13 CUB domains cause abnormal pre‐mRNA splicing and defective secretion of ADAMTS13. J Cell Mol Med 24, 4356–4361 (2020).

24.Wang, J. & Zhao, L. Clinical Features and Gene Mutation Analysis of Congenital Thrombotic Thrombocytopenic Purpura in Neonates. Frontiers Pediatrics 8, 546248 (2020).

25.Dai, Y. et al. Hereditary Thrombotic Thrombocytopenic Purpura in a Chinese Boy With a Novel Compound Heterozygous Mutation of the ADAMTS13 Gene. Frontiers Pediatrics 8, 554 (2020).

26.Das, D. et al. Novel homozygous ADAMTS13 mutation with neonatal presentation of congenital thrombotic thrombocytopenic purpura in two Indian origin babies. Pediatric Hematology Oncol J 4, 27–29 (2019).

27.Roose, E. et al. Anti-ADAMTS13 Antibodies and a Novel Heterozygous p.R1177Q Mutation in a Case of Pregnancy-Onset Immune-Mediated Thrombotic Thrombocytopenic Purpura. Th Open 02, e8–e15 (2018).

28.Schelpe, A. et al. Child‐onset thrombotic thrombocytopenic purpura caused by p.R498C and p.G259PfsX133 mutations in ADAMTS13. Eur J Haematol 101, 191–199 (2018).

29.Peyvandi, F. et al. Mechanisms of the interaction between two ADAMTS13 gene mutations leading to severe deficiency of enzymatic activity. Hum Mutat 27, 330–336 (2006).

30.Fan, X. et al. Genetic variations in complement factors in patients with congenital thrombotic thrombocytopenic purpura with renal insufficiency. Int J Hematol 103, 283–291 (2016).

31.Ferrari, B. et al. Risk of diagnostic delay in congenital thrombotic thrombocytopenic purpura. J Thromb Haemost 17, 666–669 (2019).

32.Ogawa, Y. et al. A Unique Case Involving a Female Patient with Upshaw-Schulman Syndrome: Low Titers of Antibodies against ADAMTS13 prior to Pregnancy Disappeared after Successful Delivery. Transfus Med Hemoth 42, 59–63 (2015).

33.Ferrari, B. et al. Congenital and acquired ADAMTS13 deficiency: Two mechanisms, one patient. J Clin Apheresis 30, 252–256 (2015).

34.Itami, H. et al. Complement activation associated with ADAMTS13 deficiency may contribute to the characteristic glomerular manifestations in Upshaw-Schulman syndrome. Thromb Res 170, 148–155 (2018).

35.Yadav, R. et al. Novel Heterozygous Mutations of Congenital Thrombotic Thrombocytopenic Purpura: A Rare Case Report. Indian J Nephrol 29, 295 (2019).

36.Falter, T. et al. Late onset and pregnancy-induced congenital thrombotic thrombocytopenic purpura. Hämostaseologie 34, 244–248 (2014).

37.Scully, M. et al. Thrombotic thrombocytopenic purpura and pregnancy: presentation, management, and subsequent pregnancy outcomes. Blood 124, 211–9 (2014).

38.Kovarova, P., Hrdlickova, R., Blahutova, S. & Cermakova, Z. ADAMTS13 kinetics after therapeutic plasma exchange and plasma infusion in patients with Upshaw-Schulman syndrome. J Clin Apheresis 34, 13–20 (2018).

39.Eura, Y. et al. Candidate gene analysis using genomic quantitative PCR: identification of ADAMTS13 large deletions in two patients with Upshaw-Schulman syndrome. Mol Genetics Genom Medicine 2, 240–4 (2014).

40.Rurali, E. et al. ADAMTS13 Secretion and Residual Activity among Patients with Congenital Thrombotic Thrombocytopenic Purpura with and without Renal Impairment. Clin J Am Soc Nephrol Cjasn 10, 2002–12 (2015).

41.Tenison, E., Asif, A. & Sheridan, M. Congenital thrombotic thrombocytopenic purpura presenting in adulthood with recurrent cerebrovascular events. Bmj Case Reports 12, e229481 (2019).

42.Fattah, H. et al. Successful kidney transplantation in a patient with congenital thrombotic thrombocytopenic purpura (Upshaw-Schulman syndrome): KIDNEY TRANSPLANTATION IN CONGENITAL TTP. Transfusion 57, 3058–3062 (2017).

43.Tsujii, N. et al. Severe Hemolysis and Pulmonary Hypertension in a Neonate With Upshaw-Schulman Syndrome. Pediatrics 138, e20161565–e20161565 (2016).

44.Raval, J. S., Padmanabhan, A., Hovinga, J. A. K. & Kiss, J. E. Development of a clinically significant ADAMTS13 inhibitor in a patient with hereditary thrombotic thrombocytopenic purpura. Am J Hematol 90, E22 (2014).

45.Krogh, A.-S. von et al. The impact of congenital thrombotic thrombocytopenic purpura on pregnancy complications. Thromb Haemostasis 111, 1180–1183 (2014).

46.Tanaka, H. et al. Case of maternal and fetal deaths due to severe congenital thrombotic thrombocytopenic purpura (Upshaw-Schulman syndrome) during pregnancy: Upshaw-Schulman syndrome with pregnancy. J Obstet Gynaecol Re 40, 247–249 (2013).

47.Doi, T. et al. Limited renal prophylaxis in regular plasmatherapy for heritable ADAMTS13 deficiency. Pediatr Blood Cancer 60, 1557–8 (2013).

48.Metin, A. et al. Congenital thrombotic thrombocytopenic purpura with novel mutations in three unrelated Turkish children. Pediatr Blood Cancer 61, 558–61 (2013).

49.Sissy, M. H. E., Hafez, A. A. E. & Sissy, A. H. E. Low Incidence of ADAMTS13 Missense Mutation R1060W in Adult Egyptian Patients with Thrombotic Thrombocytopenic Purpura. Acta Haematol-basel 132, 30–35 (2013).

50.Moatti-Cohen, M. et al. Unexpected frequency of Upshaw-Schulman syndrome in pregnancy-onset thrombotic thrombocytopenic purpura. Blood 119, 5888–97 (2012).

51.Lotta, L. A. et al. Residual plasmatic activity of ADAMTS13 is correlated with phenotype severity in congenital thrombotic thrombocytopenic purpura. Blood 120, 440–8 (2012).

52.He, Y., Chen, Y., Zhao, Y., Zhang, Y. & Yang, W. Clinical study on five cases of thrombotic thrombocytopenic purpura complicating pregnancy. Australian New Zealand J Obstetrics Gynaecol 50, 519–22 (2010).

53.Klukowska, A., Niewiadomska, E., Budde, U., Oyen, F. & Schneppenheim, R. Difficulties in Diagnosing Congenital Thrombotic Thrombocytopenic Purpura. J Pediatric Hematology Oncol 32, 103–107 (2010).

54.Mariotte, E. et al. Epidemiology and pathophysiology of adulthood-onset thrombotic microangiopathy with severe ADAMTS13 deficiency (thrombotic thrombocytopenic purpura): a cross-sectional analysis of the French national registry for thrombotic microangiopathy. Lancet Haematol 3, e237–e245 (2016).

55.Zharnest, D. et al. Atypical presentation of congenital thrombotic thrombocytopenic purpura with large and small vessel disease: A case report. Pediatr Blood Cancer 67, e28316 (2020).

56.Lv, H. et al. Neonate with Congenital Thrombotic Thrombocytopenic Purpura: a Case Report of a de novo Compound Heterozygote Mutation in ADAMTS13 Gene and Review of Literature. Clin Lab 66, (2020).

57.Soffer, M. D. et al. Congenital thrombotic thrombocytopenic purpura (TTP) with placental abruption despite maternal improvement: a case report. Bmc Pregnancy Childb 20, 365 (2020).

58.Pikovsky, O. et al. Congenital thrombotic thrombocytopenic purpura in a large cohort of patients carrying a novel mutation in ADAMTS13 gene. Thromb Res 185, 167–170 (2020).

59.Elbaz, C., Sholzberg, M., Hanif, H., Bonnefoy, A. & Pavenski, K. New mutation found to cause hereditary thrombotic thrombocytopenic purpura in a patient presenting with seizures in adulthood. Platelets 1–3 (2020) doi:10.1080/09537104.2020.1732327.

60.Taylor, A., Vendramin, C., Oosterholt, S., Pasqua, O. D. & Scully, M. Pharmacokinetics of plasma infusion in congenital thrombotic thrombocytopenic purpura. J Thromb Haemost 17, 88–98 (2018).

61.Beauvais, D. et al. Inherited Thrombotic Thrombocytopenic Purpura Revealed by Recurrent Strokes in a Male Adult: Case Report and Literature Review. J Stroke Cerebrovasc Dis 28, 1537–1539 (2019).

62.Alwan, F. et al. Characterization and treatment of congenital thrombotic thrombocytopenic purpura. Blood 133, 1644–1651 (2019).

63.Schneppenheim, R. et al. A common origin of the 4143insA ADAMTS13 mutation. Thromb Haemostasis 96, 3–6 (2006).

64.Prestidge, T. D. et al. Congenital thrombotic thrombocytopenic purpura (cTTP) with two novel mutations. Pediatr Blood Cancer 59, 1296–1298 (2012).

65.Peyvandi, F., Ferrari, S., Lavoretano, S., Canciani, M. T. & Mannucci, P. M. von Willebrand factor cleaving protease (ADAMTS-13) and ADAMTS-13 neutralizing autoantibodies in 100 patients with thrombotic thrombocytopenic purpura. Brit J Haematol 127, 433–439 (2004).

66.Wyatt, K. D. et al. Hereditary Thrombotic Thrombocytopenic Purpura in a 9-Month Old: Diagnosing and Managing an Ultra-rare Disorder. J Pediatric Hematology Oncol Publish Ahead of Print, (2020).

67.Ishizashi, H. et al. Quantitative Western blot analysis of plasma ADAMTS13 antigen in patients with Upshaw-Schulman syndrome. Thromb Res 120, 381–386 (2007).

68.Tanabe, S. et al. Two newborn-onset patients of Upshaw–Schulman syndrome with distinct subsequent clinical courses. Int J Hematol 96, 789–797 (2012).

69.Taguchi, F. et al. The homozygous p.C1024R- ADAMTS13 gene mutation links to a late-onset phenotype of Upshaw-Schulman syndrome in Japan. Thromb Haemostasis 107, 1003–5 (2012).

70.Deal, T., Hovinga, J. A. K., Marques, M. B. & Adamski, J. Novel ADAMTS13 mutations in an obstetric patient with Upshaw-Schulman syndrome. J Clin Apheresis 28, 311–6 (2012).

71.Pérez-Rodríguez, A. et al. A novel mutation in ADAMTS13 of a child with Upshaw-Schulman Syndrome. Thromb Haemostasis 112, 1065–8 (2014).

72.Alsultan, A., Jarrar, M., Al-Harbi, T. & Balwi, M. A. L. Novel frameshift mutations in ADAMTS13 in two families with hereditary thrombotic thrombocytopenic purpura. Pediatr Blood Cancer 60, 1559–60 (2013).

73.Lotta, L. A., Wu, H. M., Cairo, A., Bentivoglio, G. & Peyvandi, F. Drop of residual plasmatic activity of ADAMTS13 to undetectable levels during acute disease in a patient with adult-onset congenital thrombotic thrombocytopenic purpura. Blood Cells Mol Dis 50, 59–60 (2013).

74.Kim, B. et al. Single-nucleotide variations defining previously unreported ADAMTS13 haplotypes are associated with differential expression and activity of the VWF-cleaving protease in a Salvadoran congenital thrombotic thrombocytopenic purpura family. Brit J Haematol 165, 154–8 (2014).

75.CAMILLERI, R. S. et al. Prevalence of the ADAMTS-13 missense mutation R1060W in late onset adult thrombotic thrombocytopenic purpura. J Thromb Haemost 6, 331–338 (2008).

76.Snider, C. E. et al. Dissociation between the level of von Willebrand factor-cleaving protease activity and disease in a patient with congenital thrombotic thrombocytopenic purpura. Am J Hematol 77, 387–390 (2004).

77.Feys, H. B. et al. Mutation of the H-bond acceptor S119 in the ADAMTS13 metalloprotease domain reduces secretion and substrate turnover in a patient with congenital thrombotic thrombocytopenic purpura. Blood 114, 4749–4752 (2009).

78.Camilleri, R. S. et al. A phenotype-genotype correlation of ADAMTS13 mutations in congenital thrombotic thrombocytopenic purpura patients treated in the United Kingdom. J Thrombosis Haemostasis Jth 10, 1792–801 (2012).

79.Kentouche, K. et al. Remission of thrombotic thrombocytopenic purpura in a patient with compound heterozygous deficiency of von Willebrand factor‐cleaving protease by infusion of solvent/detergent plasma. Acta Paediatr 91, 1056–1059 (2002).

80.Lee, S. H. et al. A novel homozygous missense ADAMTS13 mutation Y658C in a patient with recurrent thrombotic thrombocytopenic purpura. Ann Clin Lab Sci 41, 273–6 (2011).

81.Calderazzo, J. C. et al. A new ADAMTS13 missense mutation (D1362V) in thrombotic thrombocytopenic purpura diagnosed during pregnancy. Thromb Haemostasis 108, 401–403 (2012).

82.Mise, K. et al. Long term follow up of congenital thrombotic thrombocytopenic purpura (Upshaw-Schulman syndrome) on hemodialysis for 19 years: a case report. Bmc Nephrol 14, 156 (2013).

83.Krogh, A. S. von et al. High prevalence of hereditary thrombotic thrombocytopenic purpura in central Norway: from clinical observation to evidence. J Thrombosis Haemostasis Jth 14, 73–82 (2016).

84.Pimanda, J. E. et al. Congenital thrombotic thrombocytopenic purpura in association with a mutation in the second CUB domain of ADAMTS13. Blood 103, 627–629 (2004).

85.Noris, M. et al. Complement Factor H Mutation in Familial Thrombotic Thrombocytopenic Purpura with ADAMTS13 Deficiency and Renal Involvement. J Am Soc Nephrol 16, 1177–1183 (2005).

86.Park, H. W. et al. Congenital thrombotic thrombocytopenic purpura associated with unilateral moyamoya disease. Pediatric Nephrol Berlin Ger 23, 1555–8 (2008).

87.KOKAME, K., AOYAMA, Y., MATSUMOTO, M., FUJIMURA, Y. & MIYATA, T. Inherited and de novo mutations of ADAMTS13 in a patient with Upshaw-Schulman syndrome: Letters to the Editor. J Thromb Haemost 6, 213–215 (2007).

88.Meyer, S. C. et al. A first case of congenital TTP on the African continent due to a new homozygous mutation in the catalytic domain of ADAMTS13. Ann Hematol 87, 663–666 (2008).

89.Bestetti, G. et al. ADAMTS 13 genotype and vWF protease activity in an Italian family with TTP. Thromb Haemostasis 90, 955–956 (2003).

90.Licht, C. et al. Two novel ADAMTS13 gene mutations in thrombotic thrombocytopenic purpura/hemolytic-uremic syndrome (TTP/HUS). Kidney Int 66, 955–958 (2004).

91.Shibagaki, Y. et al. Novel compound heterozygote mutations (H234Q/R1206X) of the ADAMTS13 gene in an adult patient with Upshaw–Schulman syndrome showing predominant episodes of repeated acute renal failure. Nephrol Dial Transpl 21, 1289–1292 (2006).

92.Kokame, K., Kokubo, Y. & Miyata, T. Polymorphisms and mutations of ADAMTS13 in the Japanese population and estimation of the number of patients with Upshaw-Schulman syndrome. J Thrombosis Haemostasis Jth 9, 1654–6 (2011).

93.Fujimura, Y. et al. Pregnancy-induced thrombocytopenia and TTP, and the risk of fetal death, in Upshaw-Schulman syndrome: a series of 15 pregnancies in 9 genotyped patients. Brit J Haematol 144, 742–54 (2008).

94.Donadelli, R. et al. In-vitro and in-vivo consequences of mutations in the von Willebrand factor cleaving protease ADAMTS13 in thrombotic thrombocytopenic purpura. Thromb Haemostasis 96, 454–464 (2006).

95.Palla, R. et al. The first deletion mutation in the TSP1-6 repeat domain of ADAMTS13 in a family with inherited thrombotic thrombocytopenic purpura. Haematologica 94, 289–93 (2008).

96.TAO, Z. et al. Novel ADAMTS-13 mutations in an adult with delayed onset thrombotic thrombocytopenic purpura. J Thromb Haemost 4, 1931–1935 (2006).

97.Assink, K. et al. Mutation analysis and clinical implications of von Willebrand factor–cleaving protease deficiency. Kidney Int 63, 1995–1999 (2003).

98.Garagiola, I. et al. Nonsense-mediated mRNA decay in the ADAMTS13 gene caused by a 29-nucleotide deletion. Haematologica 93, 1678–1685 (2008).

99.Studt, J.-D. et al. Fatal congenital thrombotic thrombocytopenic purpura with apparent ADAMTS13 inhibitor: in vitro inhibition of ADAMTS13 activity by hemoglobin. Blood 105, 542–544 (2005).

100.Lancellotti, S. et al. The D173G mutation in ADAMTS-13 causes a severe form of congenital thrombotic thrombocytopenic purpura: A clinical, biochemical and in silico study. Thromb Haemostasis 115, 51–62 (2016).

101.Cock, E. D. et al. The novel ADAMTS13-p.D187H mutation impairs ADAMTS13 activity and secretion and contributes to thrombotic thrombocytopenic purpura in mice. J Thromb Haemost 13, 283–292 (2015).

102.Kim, H.-Y. et al. Congenital thrombotic thrombocytopenic purpura (Upshaw-Schulman syndrome) caused by novel ADAMTS13 mutations. Brit J Haematol 173, 156–159 (2015).

103.Krabbe, J. G. et al. Adult-onset congenital thrombotic thrombocytopenic purpura caused by a novel compound heterozygous mutation of the ADAMTS13 gene. Int J Hematol 102, 477–481 (2015).

104.Epperla, N., Hemauer, K., Friedman, K. D., George, J. N. & Foy, P. Congenital thrombotic thrombocytopenic purpura related to a novel mutation in ADAMTS13 gene and management during pregnancy. Am J Hematol 91, 644–646 (2016).

105.Underwood, M., Peyvandi, F., Garagiola, I., Machin, S. & Mackie, I. Degradation of two novel congenital TTP ADAMTS13 mutants by the cell proteasome prevents ADAMTS13 secretion. Thromb Res 147, 16–23 (2016).

106.Conboy, E. et al. A Severe Case of Congenital Thrombotic Thrombocytopenia Purpura Resulting From Compound Heterozygosity Involving a Novel ADAMTS13 Pathogenic Variant. J Pediatric Hematology Oncol 40, 60–62 (2018).

107.Yadav, S., Shetty, S. & Kulkarni, B. A novel homozygous frameshift mutation in Exon 7 of the ADAMTS13 gene in a patient with congenital thrombotic thrombocytopenic purpura from India: a case report: NOVEL MUTATION CAUSING CONGENITAL TTP. Transfusion 57, 2712–2714 (2017).

108.Sarmiento, H. et al. Upshaw-Schulman Syndrome: Novel homozygous missense mutation. Thromb Res 158, 83–85 (2017).

109.Hassenpflug, W. et al. Genetic and Functional Characterization of ADAMTS13 Variants in a Patient Cohort with Upshaw–Schulman Syndrome Investigated in Germany. Thromb Haemostasis 47, 709–722 (2018).

110.Joly, B. S. et al. ADAMTS13 Gene Mutations Influence ADAMTS13 Conformation and Disease Age-Onset in the French Cohort of Upshaw–Schulman Syndrome. Thromb Haemostasis 118, 1902–1917 (2018).

111.Resham, S., Fadoo, Z. & Moiz, B. Upshaw-Schulman Syndrome With c.2728C>T Mutation in ADAMTS13 Gene. J Pediatric Hematology Oncol 41, e60–e62 (2019).

112.Rossio, R. et al. Two novel heterozygote missense mutations of the ADAMTS13 gene in a child with recurrent thrombotic thrombocytopenic purpura. Blood Transfus Trasfusione Del Sangue 11, 241–4 (2012).

113.Rank, C. U. et al. Congenital thrombotic thrombocytopenic purpura caused by new compound heterozygous mutations of the ADAMTS13 gene. Eur J Haematol 92, 168–171 (2013).

114.Hou, L. & Du, Y. Two novel mutations in ADAMTS13 in a Chinese boy with congenital thrombocytopenic purpura: a case report. Bmc Med Genet 21, 57 (2020).

115.Dorland, H. A. van et al. The International Hereditary Thrombotic Thrombocytopenic Purpura Registry: key findings at enrollment until 2017. Haematologica 104, 2107–2115 (2019).

116.Schneppenheim, R. et al. von Willebrand factor cleaving protease and ADAMTS13 mutations in childhood TTP. Blood 101, 1845–1850 (2003).

117.Matsumoto, M. et al. Molecular characterization of ADAMTS13 gene mutations in Japanese patients with Upshaw-Schulman syndrome. Blood 103, 1305–1310 (2004).

118.Uchida, T. et al. Identification of novel mutations in ADAMTS13 in an adult patient with congenital thrombotic thrombocytopenic purpura. Blood 104, 2081–2083 (2004).

119.Savasan, S., Lee, S.-K., Ginsburg, D. & Tsai, H.-M. ADAMTS13 gene mutation in congenital thrombotic thrombocytopenic purpura with previously reported normal VWF cleaving protease activity. Blood 101, 4449–4451 (2003).

120.Kokame, K. et al. Mutations and common polymorphisms in ADAMTS13 gene responsible for von Willebrand factor-cleaving protease activity. Proc National Acad Sci 99, 11902–11907 (2002).

121.Levy, G. G. et al. Mutations in a member of the ADAMTS gene family cause thrombotic thrombocytopenic purpura. Nature 413, 488–494 (2001).

122.Plaimauer, B. et al. Modulation of ADAMTS13 secretion and specific activity by a combination of common amino acid polymorphisms and a missense mutation. Blood 107, 118–125 (2006).

123.Antoine, G. et al. ADAMTS13 gene defects in two brothers with constitutional thrombotic thrombocytopenic purpura and normalization of von Willebrand factor-cleaving protease activity by recombinant human ADAMTS13. Brit J Haematol 120, 821–824 (2003).

124.Veyradier, A. et al. Ten candidate ADAMTS13 mutations in six French families with congenital thrombotic thrombocytopenic purpura (Upshaw-Schulman syndrome). J Thromb Haemost 2, 424–429 (2004).

125.Hommais, A. et al. Molecular characterization of four ADAMTS13 mutations responsible for congenital thrombotic thrombocytopenic purpura (Upshaw-Schulman syndrome). Thromb Haemostasis 98, 593–599 (2007).

126.Fujimura, Y. et al. Natural history of Upshaw-Schulman syndrome based on ADAMTS13 gene analysis in Japan. J Thrombosis Haemostasis Jth 9 Suppl 1, 283–301 (2011).

### Supplementary Tables

**Known sheet**: all ADAMTS13 variants mined from PubMed and Scopus literature.

**KnownUnique sheet**: all ADAMTS13 reported variants.

**All sheet**: all database and reported variants collection from ClinVar, HGMD with gnomAD allele frequency.

**AllUnique sheet**: all ADAMTS13 variants collection

**AllUniqueLite sheet**: all ADAMTS13 variants collection with gnomAD allele frequency.

**Data sheet**: eight population ADAMTS13 genetic prevalence and carrier frequency.

**ClinVar sheet**: ADAMTS13 variants in ClinVar database

**gnomAD sheet**: ADAMTS13 variants in gnomAD database

**HGMD sheet**: ADAMTS13 variants in HGMD database
